# Influence of mould growth and outdoor exposure on the efficacy of attractive targeted sugar baits in western kenya

**DOI:** 10.1101/2024.11.28.625847

**Authors:** Nick Yalla, Jackline Kosgei, Frank Mechan, Daniel P. McDermott, Brian Polo, Seline Omondi, Elizabeth Omukunda, Eric Ochomo

## Abstract

**Introduction:** Attractive targeted sugar baits (ATSBs) are effective against *Anopheles* mosquitoes in semiarid climates with low humidity. High humidity, however, promotes growth of moulds on the surface of ATSBs. The impact of mould on ATSB efficacy against malaria vectors remains unknown. This study explored how mould growth affects the performance of ATSB version 1.2 by comparing mouldy stations from exposed environments to non-mouldy stations from protected settings through laboratory bioassays with the local malaria vector, *Anopheles arabiensis*.

**Methods:** One hundred ATSB stations were deployed in Asembo, Rarieda-Subcounty, Siaya County, with six samples (three mouldy from exposed locations and three non-mouldy from protected locations) collected monthly for laboratory bioassays. These were tested alongside three new laboratory-kept ATSBs and two negative controls (water only and 77% sugar solution with water) to assess mosquito feeding and mortality over 48 hours.

**Results:** This study found that after 12 months of outdoor exposure, the mouldiest ATSBs from exposed locations showed a non-significant reduction in *Anopheles arabiensis* feeding rates compared to the least mouldy ATSBs from protected locations 57.42% (95% CI: 45.64-68.85) vs. 74.40% (95% CI: 64.56-82.50), respectively (P =0.062). Mosquito mortality significantly declined on mouldy ATSBs compared to laboratory controls (95% CI: 92.23-97.48) vs. 98.70% (95% CI: 97.87-99.30) respectively (P = 0.002). In contrast, protected (non-mouldy) ATSBs showed only a slight reduction in mortality compared to controls 95.94% (95% CI: 90.42-97.46) vs. 98.91% (95% CI: 97.67-99.60) respectively (P = 0.009).

**Conclusion:** This study provides evidence that environmental exposure post-deployment slightly reduced the efficacy of ATSBs in controlling *Anopheles arabiensis*, particularly beyond the recommended 6-month period. Although mould may have contributed to this reduction over 12 months, no significant difference was found between mouldy and non-mouldy ATSBs. However, mould invasion and community concerns highlight the need to replace mouldy stations to maintain effectiveness and safety.

## Background

Attractive Targeted Sugar Bait (ATSB) is a novel outdoor tool that exploits mosquito sugar-feeding behaviour to control them [1]. The concept of ATSB allows a range of insecticides or toxicants to be delivered to the mosquitoes through ingestion [2]. This has led to explorations with multiple insecticides in mosquito control. Chemical classes of insecticides that have already been tested as ATSBs include pyrethroids, carbamates, organophosphates, ivermectins, borates, neonicotinoids, spinosyns, pyriproxyfen, pyrroles, double-stranded RNA (dsRNA), biopesticides and phenylprazoles [1-9].

Previously, ATSB was implemented by spraying non-flowering vegetation close to mosquito larval habitats, targeting the newly emerged sugar-seeking vectors [10]. This approach notably decimated mosquito populations of different genera in varied study regions [11, 12] but could be environmentally unsustainable because of the potentially hazardous impact on non-target organisms (NTOs).

The version 1.2 ATSB stations used in this experiment contained fruit syrup as an attractant and stimulant for inducing the mosquitoes to feed and ingest a lethal dose of dinotefuran, a neonicotinoid insecticide, and were manufactured by Westham Co. Hod-Hasharon, Israel. They also contained uranine as a fluorescent tracer. This ATSB is designed to be hung outdoors on a wall under a roof to control mosquitoes in the peridomestic space [13]. A previous version of this ATSB station was tested in Mali and achieved a significant density reduction of 90% of female *Anopheles gambiae s*.*l*. past their second gonotrophic cycle [14].

When the new version 1.2 of the same ATSB product was deployed in western Kenya for a large-scale epidemiological trial, mould growth was observed on the many bait station membranes. This growth was observed through regular monitoring to be widespread on structures with short roof lengths overhanging the walls where the ATSBs were mounted, so the products were exposed to weather elements like sun and rainfall. In Zambia, mould growth contributed to approximately 28% of all ATSB damage, and it was closely linked to the duration of deployment. ATSBs that sustained tears were particularly vulnerable to mould attacks, especially when left in the field for long periods without replacement. Furthermore, the characteristics of the deployment site, for example whether the ATSB was protected from rainfall or not, also played a role in predicting the likelihood of mould damage [15].

Most moulds favourably proliferate on paper products, cardboard ceiling tiles, wood products, insulation materials, upholstery, and other fabrics, breaking them down [16]. The ideal growth conditions for these moulds are moisture availability, carbohydrates like sucrose, and sufficient oxygen. Low diurnal temperature ranges and high humidity characterize the western Kenya region, particularly during the long rainy season. Moreover, ATSB is approximately 70% sugar solution, a carbohydrate readily broken down by the mould. This provides favourable conditions for mould growth on the bait station membranes that might be expected to reduce the efficacy of ATSBs in controlling malaria vectors by either lowering their attractiveness to or feeding rates on the bait stations by the target vectors. Some moulds are toxic, while others have been useful to humans in different ways. There has been no reported work that has aimed to characterise the mould species that grow on the ATSBs in the areas where they have been deployed. The current policy is to remove the bait stations on walls if they become mouldy with mould spots bigger than a pencil eraser with a diameter of approximately 0.5 inches [13]. This replacement criterion can increase wastage and pose a challenge for intervention coverage when there isn’t a rigorous monitoring and replacement programme. Additionally, there are concerns about the impact of mould growth on people’s health.

There is currently no documented evidence of the potential impact of mould growth on ATSB’s ability to affect mosquito mortality or deter feeding. This study investigated the influence of mould growth on the efficacy of ATSB in causing mosquito mortality by comparing the feeding rates of malaria vectors on mouldy and non-mouldy ATSBs in laboratory bioassays. In addition, this study evaluated the effectiveness of the ATSBs beyond the six months recommended by the manufacturer to assess the persistence of the AI. The results of this study provided insights into whether the natural presence of the mould can impact the efficacy of ATSB as a complementary outdoor malaria vector control strategy in western Kenya.

## Methods

### Study site & design

One hundred uranine-dye stained ATSBs were hung in Asembo, which is part of the study site for the large-scale, cluster-randomized controlled trial with ATSBs implemented between March 2022 and March 2024 [13]. Fifty ATSBs were hung on a structure wall without a roof overhang, termed herein as (an exposed location), and another fifty were hung on a separate wall with a roof overhang, termed herein as (a protected location), protecting the bait stations from direct effects of weather elements such as rain, sunlight, and wind. The ATSBs were hung on the walls 1.8 meters from the ground.

### Mould culturing and morphological identification

To investigate and isolate the mould growing on ATSB stations, six bait stations that exhibited significant mould growth and were more than six months old were randomly selected from Asembo and transported to the laboratory for culturing. A pair of forceps was sterilized by dipping in 70% ethanol and subsequently briefly flamed over a Bunsen burner. The sterile forceps was used to gently scrape off a small portion of mould from the ATSB surface, ensuring that both mouldy and adjacent areas were included to capture all species of fungi growing. 4.8g of potato dextrose agar (PDA) was weighed and transferred to a sterile conical flask. The PDA was then suspended in 120 ml of distilled water, stirred for 1 minute using a magnetic stirrer, and heated on a hot plate until the medium was completely dissolved. The medium was autoclaved for 20 minutes at 121°C and 15 psi. After autoclaving, the media was allowed to cool to about 45ºC, permitting the gel to solidify. 10% tartaric acid was added to the medium to lower the pH to approximately 3.5 to inhibit bacterial growth. Subsequently, 20 ml of the medium was dispensed into six petri dishes each. The spread plate streaking technique was applied to isolate the mould inoculum, picked from six different ATSBs, onto each petri dish. The plates were then incubated for 7 days at 27ºC. Finally, the colony was observed under a light microscope (Leica DM750M, Suzhou, China) to reveal the fungal morphology.

### Mosquito samples

*Anopheles* mosquitoes used in the tests were collected as larvae from Ahero (0.16°S, 35.0°E) in Kisumu County, Kenya, using the larval dipping method. The field-collected larvae were raised under laboratory conditions with temperature maintained at 28 ± 2 ºC and a relative humidity of 80 ±10%. The larvae were fed on TetraMin fish food up to the pupal stage, after which they were transferred to plastic cups using 3ml Pasteur pipettes. The cups containing pupae were placed in (30×30×30) cm cages for emergence. Adults were maintained on a 10% sugar solution soaked in cotton wool until 3-5 days post-emergence, when they were used for bioassays. 204 of the mosquitoes were analysed by PCR for species identification.

### Mosquito-ATSB feeding rates and mortality assessment

Field ATSBs were selected monthly for bioefficacy tests in the KEMRI laboratory between October 2022 and October 2023, a time span longer than the manufacturer’s prescribed period of six months [13]. Every month, six ATSBs, three non-mouldy from the protected and three mouldy from the exposed locations, were removed and transported safely in portrait orientation in wooden transportation boxes of dimensions 35 cm x 25 cm x 7cm to the KEMRI laboratory in Kisumu County for bioassays. The most mouldy ATSBs from the exposed location and non-mouldy from the protected location were scouted and selected for bioassay. ATSBs selected for bioassay were selected if more than half of the bait station membrane had mould covering their surfaces. The protected location had a roof overhang, which provided protection to the ATSBs, which based on observations from the trial slowed the mould growth rates compared to those without roof protection. In addition, three laboratory-kept ATSBs that had never been used were tested alongside the field bait stations each month to assess *Anopheles* mosquito feeding rates in the laboratory. The laboratory kept ATSBs were from the same manufacturing batch as the field collected ones and were freshly opened from their packaging materials at the start of the experiment. The laboratory ATSB storage condition was 60% relative humidity and temperature of 25°C. The field-collected most mouldy, non-mouldy, and laboratory-kept ATSBs were tested for mosquito feeding rates in cages with *ad libitum* access to water and negative controls of water alone and water with 77% sugar solution (considered similar to ATSB in sugar concentration). All the test cages were supplied with ad libitum water including the 77% sugar cage (Fig. 1).

**Fig 1.**
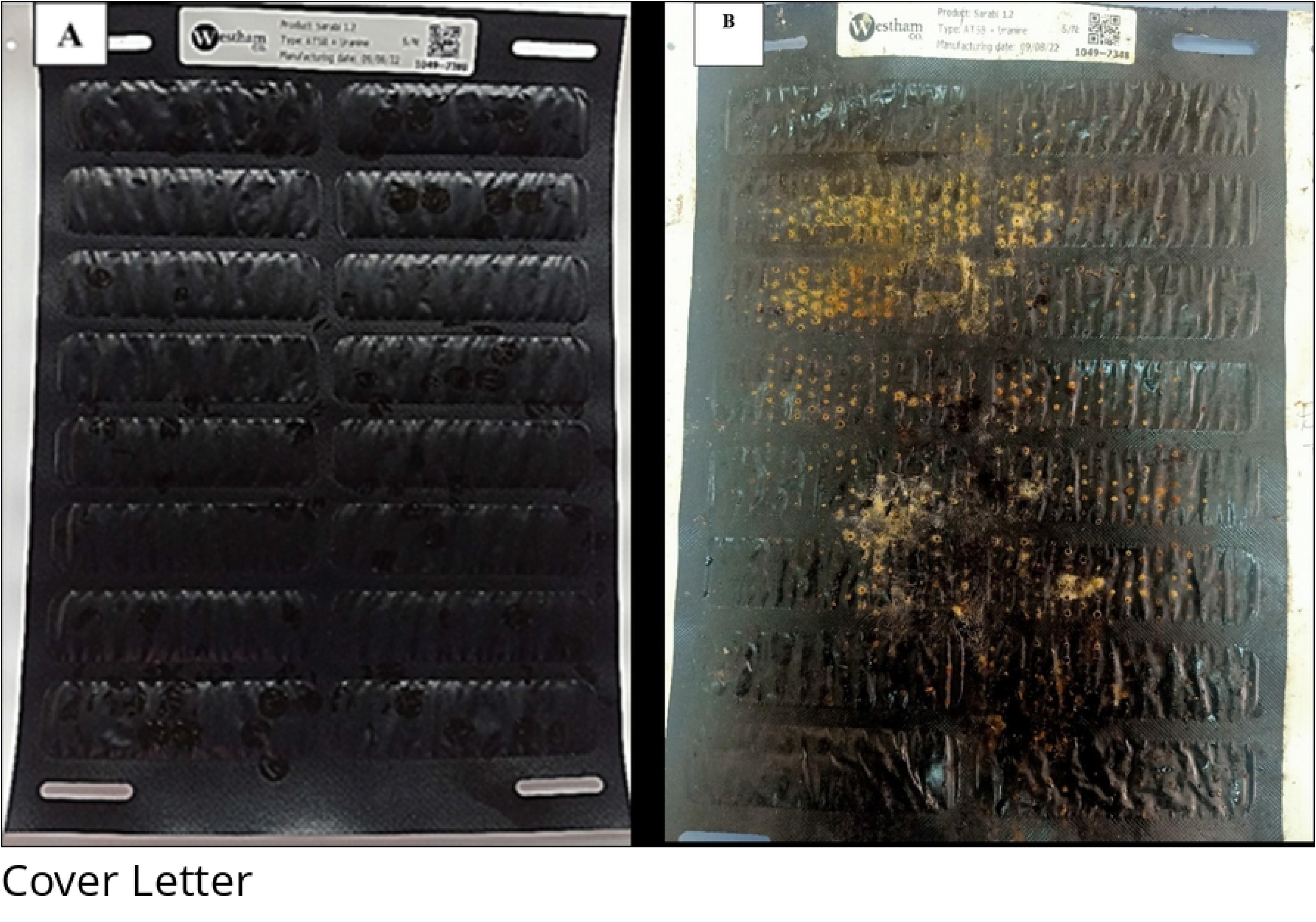
Pictures of new ATSB freshly removed from packaging laboratory kept (A) and field-collected mouldy ATSBs (B), respectively.

Mosquito bioassays were conducted in plastic Bugdorm cages measuring 30×30×30 cm. Each cage was fitted with one ATSB, positioned vertically on one of the four walls of the cage frame, and firmly secured with masking tape. Including the controls, cohorts of 3-5 day-old mosquitoes (60 males and 60 females) were starved 24 hours prior to the experiment and released to the cages containing the bait stations. The mosquitoes were allowed to feed on the bait stations for 48 hours, starting in the early afternoon at 1500 hours. At the 24-hour time point, all dead male and female mosquitoes in the treatment and control cages were aspirated out of the cages and placed in specimen vials. The remaining live mosquitoes were left in the cages with the bait station and monitored for a further 24 hours. At the end of 48 hours, all dead and alive mosquitoes in all treatments and control cages were aspirated out. The live mosquitoes were killed by freezing in a -20°C freezer. All the samples, including those sampled at 24 hours, were examined for uranine dye to detect if they had fed on the ATSB. This was done by scanning through the mosquito abdomens with a fluorescent microscope (Model: Leica Mz 10f,10 Parkway North Blvd, Suite 300, Deerfield, Illinois 60015 United States). The bio-assayed mosquitoes were individually placed onto the fluorescent microscope stage in a Petri dish directly under the field of view to assess the presence of uranine dye. White light was used for microscopic illumination, and a green UV filter was used to visualize uranine fluorescence. An observation of a light green colour of uranine dye in the mosquito’s abdomen/gut indicated feeding, while the absence of the dye colour meant an unfed mosquito.

### Statistical analysis

Generalized Linear Mixed Models (GLMMs) in R statistical software analyzed associations between station location, time-point, and outcomes (proportion fed and proportion dead). A random effect term was included to account for unexplained variation between assays identified by the ID of each bait station. ATSB location was treated as a categorical variable, while the time-point was considered a continuous variable. The statistical significance of each parameter, including an interaction term between station location and time-point, was evaluated using log-likelihood ratio tests (LRTs), comparing models with and without the parameter. Post hoc analysis was conducted using the ‘lsmeans’ package.

### Ethical considerations

The study was approved by the Kenya Medical Research Institute/Scientific and Ethics Review Unit (KEMRI/SERU), number (SERU 3613). It was also approved by the Institutional Review Board of the US Centres for Disease Control and Prevention (#7112) and the Liverpool School of Tropical Medicine Research Ethics Committee (18-015) in agreement with KEMRI SERU.

## Results

### Mould culture identification

The PDA culture characteristics of the mould revealed a mixed colony of fungal species that initially appeared cottony white after two days of incubation but turned black in larger parts of the petri dishes after seven days, producing conidial spores. By the end of the incubation period, most of the culture remained black. The fungal culture was examined under a light microscope (Leica DM750M, Suzhou, China). *Aspergillus niger* mould was identified to be the dominant species in the culture medium in all the plates assessed (Fig. 2).

**Fig 2:**
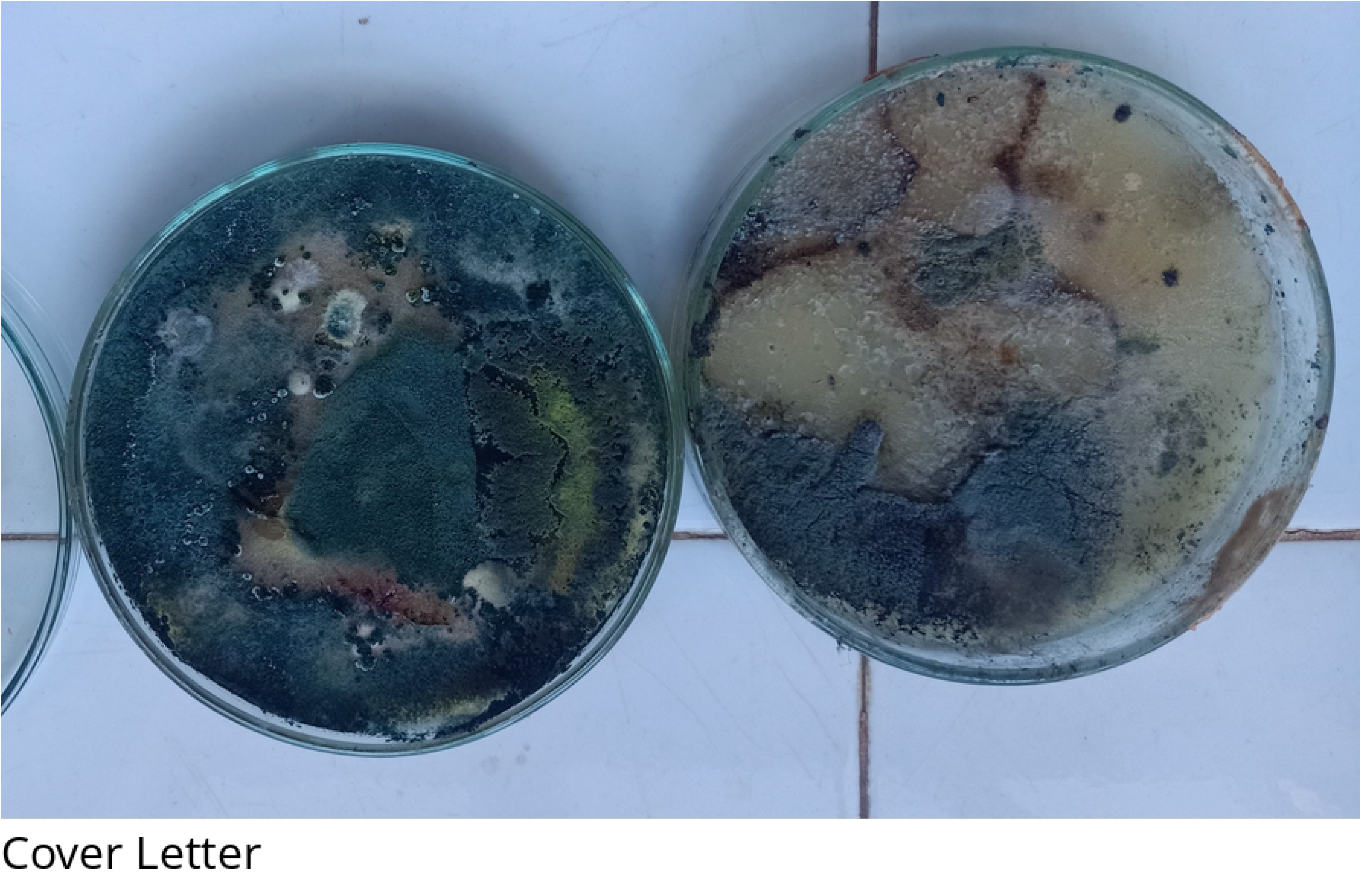
Picture of mould colony cultured on potato dextrose agar on day seven.

### Mosquito samples

Ninety-five percent of the 204 Anopheles mosquitoes tested were confirmed to be *An. arabiensis* by PCR, while ten samples did not amplify.

### Cumulative Mosquito feeding rates

Time and bait station location (‘exposed’ mouldy vs. ‘protected’ non-mouldy) were significant predictors of the cumulative proportion of mosquitoes showing dye-positive results at 48 hours (P < 0.001 for both). A significant interaction between the time-point and bait station location (P < 0.001) was observed (Fig 3A). By 5 months, the ATSBs in the field had a significantly lower mosquito feeding rate compared to the laboratory control, with both below 90%, with rates of 87.82% (95% CI: 84.12-91.09) and 92.63% (95% CI: 90.81-94.17) respectively (P = 0.001). Owing to the importance of feeding rate to the success of the product this may present a concern to efficacy in the later part of the deployment cycle. ATSBs sampled from the protected location exhibited significantly lower dye-positivity rates compared to the lab-kept ATSBs at month five, with rates of 86.34% (95% CI: 82.78-89.11) and 92.63% (95% CI: 90.81-94.17) respectively (P < 0.001). This study did not find a significant difference between the exposed (mouldy) and protected (non-mouldy) stations at any one time point however, the mouldy stations’ feeding rate did fall at the final time point with rates of 57.42% (95% CI: 45.64-68.85) vs. 74.40% (95% CI: 64.56-82.50), respectively (P = 0.062)

**Fig 3:**
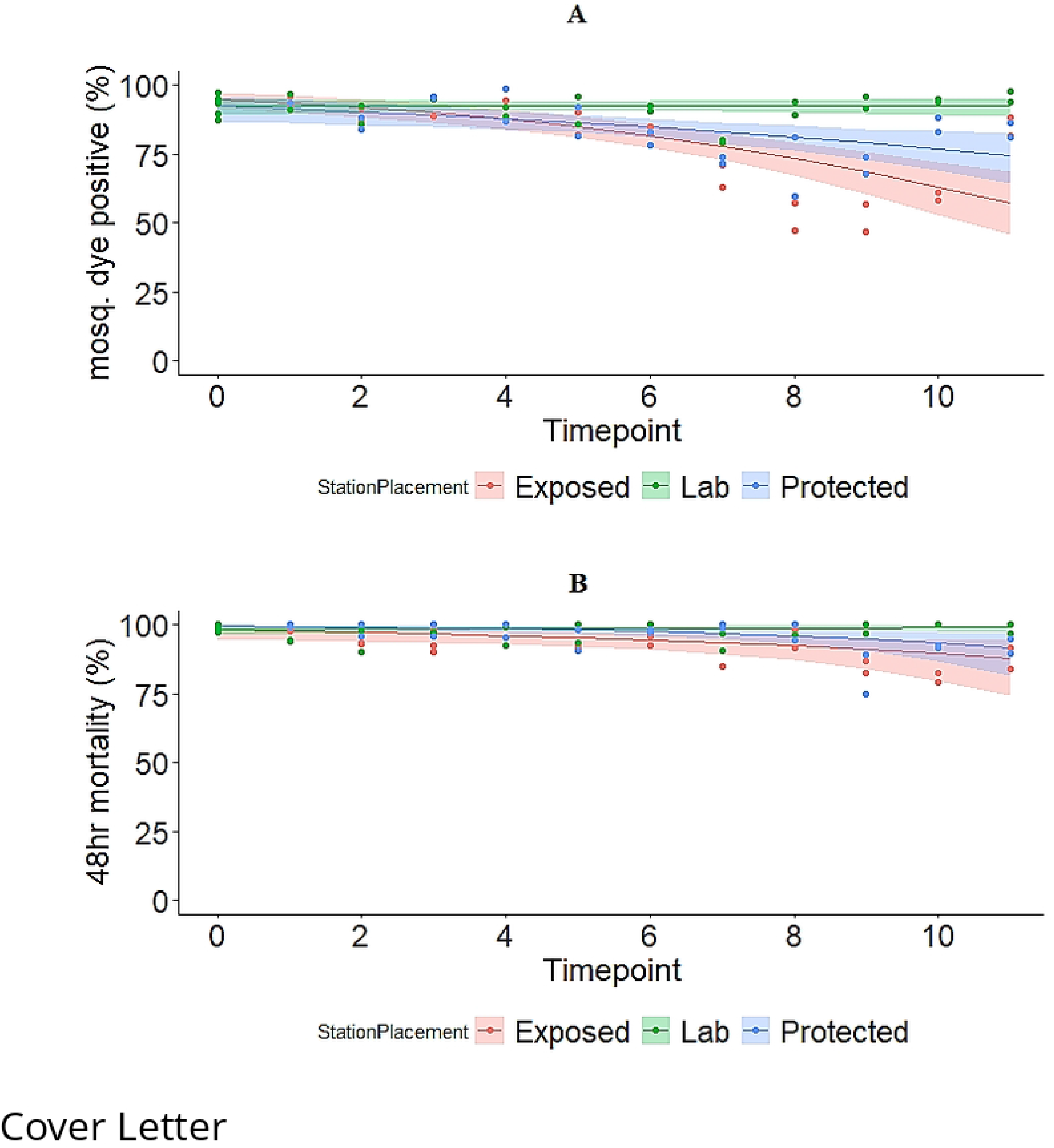
Mosquito dye positive and 48-hour mortality rates (A and B) on ATSBs, respectively. The coloured bands represent 95% intervals.

### Cumulative Mosquito Mortality rates at 48 hours

Time and bait station location were found to be significant predictors of the cumulative proportion of mosquito mortality at 48 hours (P = 0.009 and P = 0.001, respectively). A significant interaction between the time-point and bait Station location (P < 0.001) was observed (Fig 3B). The exposed location (Mouldy) did see a significant reduction in mortality compared to laboratory control by month five with rates of 95.35% (95% CI: 92.23-97.48) compared to 98.70% (95% CI: 97.87-99.30) respectively (P = 0.002). The mortality estimated dropped in month eight, which would raise concerns about its use beyond this period. The protected location experienced a small reduction in mortality by the seventh month compared to the laboratory controls, 95.94% (95% CI: 90.42-97.46) vs. 98.91% (95% CI: 97.67-99.60) respectively (P = 0.009), though the mortality rate remained above 95% until the eighth month. These findings demonstrate that, despite mould growth, the stations effectively delivered a toxic dose of the AI up to the twelfth month. However, extending the deployment period beyond the current six months without further station optimization raises concerns. There were no significant differences between the exposed and protected ATSBs at any time-point, with mean 48-hour mortality rates at the final time-point being 87.76% (95% CI: 74.17-94.37) vs. 91.54% (95% CI: 81.48-96.83) respectively (P = 0.807). Less than 10% and 5% cumulative mortalities were recorded in water only and 77% sugar solution plus water controls, respectively.

## Discussion

This study established that during the recommended lifespan, when exposed to environmental conditions that lead to mould growth, the ATSBs still had a similar feeding rate in cage experiments up to five months post-deployment and similarly high mortality of mosquitoes to the laboratory control baits. The feeding and mortality rates of those outdoors declined slowly, with feeding rates remaining >85% up to five months in both exposed and protected conditions. While no statistical difference in feeding rate was detected between exposed (mouldy) and protected (non-mouldy) at any one time point, the exposed stations had a lower feeding rate overall and the low feeding rate for mouldy ATSBs particularly after 8 months warrants further study.

Either the presence of mould or other environmental factors differing between exposed and protected environments seems to alter the mosquitoes’ ability to feed on ATSBs. The mould might be blocking the ATSB pores or hardening the membrane surfaces, making it difficult for mosquitoes to probe and ingest the toxin. This suggests that substantial mould growth on the bait station may warrant the removal or replacement of the product from the walls if better mosquito control with ATSB is to be attained in environments favouring fungal growth. Culturing revealed different fungal species growing on the product. Our study identified *Aspergillus niger* as the most dominant mould in the colony but, unfortunately, failed to characterize other mould species that formed part of the culture. Given the few samples of mouldy ATSBs assayed, this study might not have involved all the types of mould growing on the product in the entire study area, and further characterisation studies are needed. The safety level of most of these moulds that grow on the product is still unknown. Thus it remains a concern that despite not detecting a major problem for efficacy in laboratory feeding and mortality tests, the risk of exposure to these moulds needs further investigation. Mould damage to ATSB stations appears to be linked to deployment duration with an inverse relationship observed between mould prevalence and tear damage [15]. Rapid identification and replacement of torn ATSBs may therefore reduce the rate of development of mould but this might require high bait station turnover. However, frequent replacement of the ATSBs is not sustainable and might make deployment very expensive. Site characteristics and weather elements predict the likelihood of damage types [15] like mould. Mould and other types of damage, may also influence community acceptance, making it important to balance operational feasibility with community perceptions for successful deployment [15]. *Aspergillus niger*, found to be the most dominant mould in this study, is, however, not considered harmful to humans in lower doses, with between 200-500 spores typically not an issue and staying within the normal range. Moreover, when Penicillium, Aspergillus, or Cladosporium levels range between 500-1500 spores, remediation is typically not required as they are regarded not harmful to human [17-19]. Most people potentially breathe in *Aspergillus* spores every day without getting sick, as *Aspergillus niger* is generally regarded as a non-pathogenic fungus widely distributed in nature. Only in a few cases has *Aspergillus niger* has been able to colonize the human body as an opportunistic invader, and in almost all these cases, the patients have a history of severe illness or immunosuppressive treatment [20]. However, the authors recognise the need for a more rigorous screening strategy to identify if any mould that may cause pathology in household members who are in close proximity to the ATSB can be found. Even in the absence of efficacy concerns and health concerns, the community acceptability of an intervention that grows mould on people’s homes may present an issue.

Given these concerns, there is a need to alter the design or increase the anti-mould agents in the product to reduce the rate of mould development, especially given the high temperature and high humidity environments this product may eventually be deployed in. It is essential to note that weather conditions, particularly humidity, play a significant role in the propagation of mould in addition to moisture, carbohydrate substrate, and moderately low-temperature conditions [15]. Western Kenya provides conducive conditions to mould growth compared to Mali, which is drier for the better part of the year. This probably explains the lack of mould occurrence in Mali during the testing of ATSBs there as opposed to the current setting [14]. Further, this difference in climate highlights the importance of considering environmental factors when assessing the efficacy of vector control tools, especially those aimed to be deployed for outdoor use. More research is needed on how mould on ATSB impedes mosquito feeding. Whether it blocks or hardens the bait station membrane making it impossible for the mosquito proboscis to pierce and suck the bait solution, is unknown. In the natural setting, mould might significantly lower the product efficacy through reduced feeding rates due to blocked pores or hardened membrane surfaces of the ATSBs. The observed bigger reduction of mosquito feeding rate than mosquito mortality in this study was possibly due to dried active ingredient on the ATSB membrane surfaces. Future studies should consider collecting weather data to evaluate the dynamics of mosquito feeding rates as well as mould growth, with consideration of temperature, rainfall, and humidity, providing a more comprehensive understanding of the impact of weather variables on mould growth and the product’s effectiveness. This feeding assay was conducted in cages, which may not fully replicate mosquito behaviour in natural settings. With only one hundred ATSBs used and only 3 mouldy and 3 non-mouldy ATSBs compared at each time point, the study suggests that future research should include a larger sample size to provide more conclusive insights into the impact of mould on ATSB performance. Worth noting is the fact that bait station location was confounded with mould presence, making it impossible to distinguish these factors in this study. Future studies should consider using ATSBs from the same environment.

## Conclusion

This study provided evidence that environmental exposure of the Westham ATSB station v1.2 in western Kenya led to a slight but progressive reduction in the efficacy of the product in controlling *Anopheles arabiensis* mosquitoes in terms of diminishing the feeding and mortality rates in laboratory bioassays, particularly beyond the end of the product’s 6-month recommended deployment period. While there was evidence that the presence of mould exacerbated this effect over a 12-month outdoor exposure period, there was no significant efficacy difference at any one-time point between the stations that had been outdoors and were mouldy and those that had been outdoors and were not mouldy. Owing to the overall reduction seen, the absence of safety data around the moulds that are present, and community perceptions, we still recommend the removal and replacement of any ATSB stations that become torn or mouldy.

## Acknowledgements

We are grateful to the Entomology field team, which collected the samples to run the assays. This study was funded by IVCC through support from the Bill & Melinda Gates Foundation (grant INV-007509), the Swiss Agency for Development and Cooperation (SDC) (grant 81067480) and UK Aid ((grant 30041-105). The funders had no role in the study design, data collection, analysis, the decision to publish, or the preparation of the manuscript. The findings and conclusions contained within are those of the authors and do not necessarily reflect positions or policies of the Bill & Melinda Gates Foundation, SDC or UK Aid. Westham Co Ltd, Hod-Hasharon, Israel, provided ATSB samples.

## Authors’ contributions

NY conceptualization, sample collection, data collection, analysis, and manuscript drafting. JK, BP, SO, EO, and EO supervised and reviewed the manuscript. DM and FM data analysis and review of the manuscript. All authors read and approved the final manuscript.

## Supporting information

S1_Study Data.

